# MosaicPlanner: Hardware Agnostic Array Tomography Acquisition Software

**DOI:** 10.1101/473009

**Authors:** R. Serafin, O. Gliko, S. J Smith, F. Collman

**Affiliations:** Allen Institute for Brain Science, Seattle WA 98109

**Keywords:** array tomography, fluorescence microscopy, imaging software

## Abstract

Array tomography (AT) is a technique for acquiring high resolution highly multiplexed imagery from series of ultra-thin sections arranged as an array on a rigid substrate. Specialized microscope control has been required to utilize AT as an imaging technique, which is often time consuming, and yields small volume data sets. Here we present MosaicPlanner, an open source software platform for light level AT, that streamlines the acquisition process and utilizes the general microscope control API provided by Micro-Manager, allowing AT data to be acquired on a wide variety of microscope hardware. This report provides a description of the MosaicPlanner software design, and platform improvements that were implemented to increase the acquisition speed of high volume, multiplexed AT datasets.

## Introduction

In biological disciplines three-dimensional tissue visualization is paramount to research. In fluorescence microscopy, there is particular interest in not only the ability to resolve the morphological structures of a sample with high precision, but the capability to localize multiple unique molecular species expressed in the tissue within one experiment. AT is a high resolution, highly multiplexable imaging technique of ultra thin serial sections placed on a rigid substrate [*Micheva & Smith 2007*]. In addition to the degree of multiplexing with molecular specificity possible with AT [*Collman et al. 2015*] another major advantage of the technique is the sub diffraction limit axial resolution provided by the two-dimensional nature of the sample. This presents an increase in complexity for data acquisition, as individual sections are arranged across the substrate, in a semi-regular but not trivially predictable pattern. AT samples are typically sectioned at 50 - 100 nm, which inevitably means that a large number of images are required to aptly study biological morphologies. The complexity of executing an AT imaging experiment rapidly multiplies not only with increasing volume size of a sample, but also the desired degree of multiplexing within an experiment. Furthermore, keeping all these image data organized to facilitate the necessary post-acquisition image processing can be its own challenge. Here we present MosaicPlanner, a generalizable AT software platform designed to address these challenges by facilitating automated setup, execution, and analysis of AT imaging experiments.

In order to effectively set up an AT acquisition, a microscopist needs to be able to visualize their sample spatially. To utilize the full potential of AT an experimenter must set up a series of stereotyped tiling sequences across multiple sections which cover precisely the same region of tissue (Figure 1a). Further, if one wants to be able to execute multiple rounds of multiplexed imaging on the same sample the microscopist needs a reproducible way to set up and execute individual staining rounds within an imaging experiment. This ‘mapping’ problem can make setting up AT imaging experiments a tedious and time consuming process.

**Figure 1.**
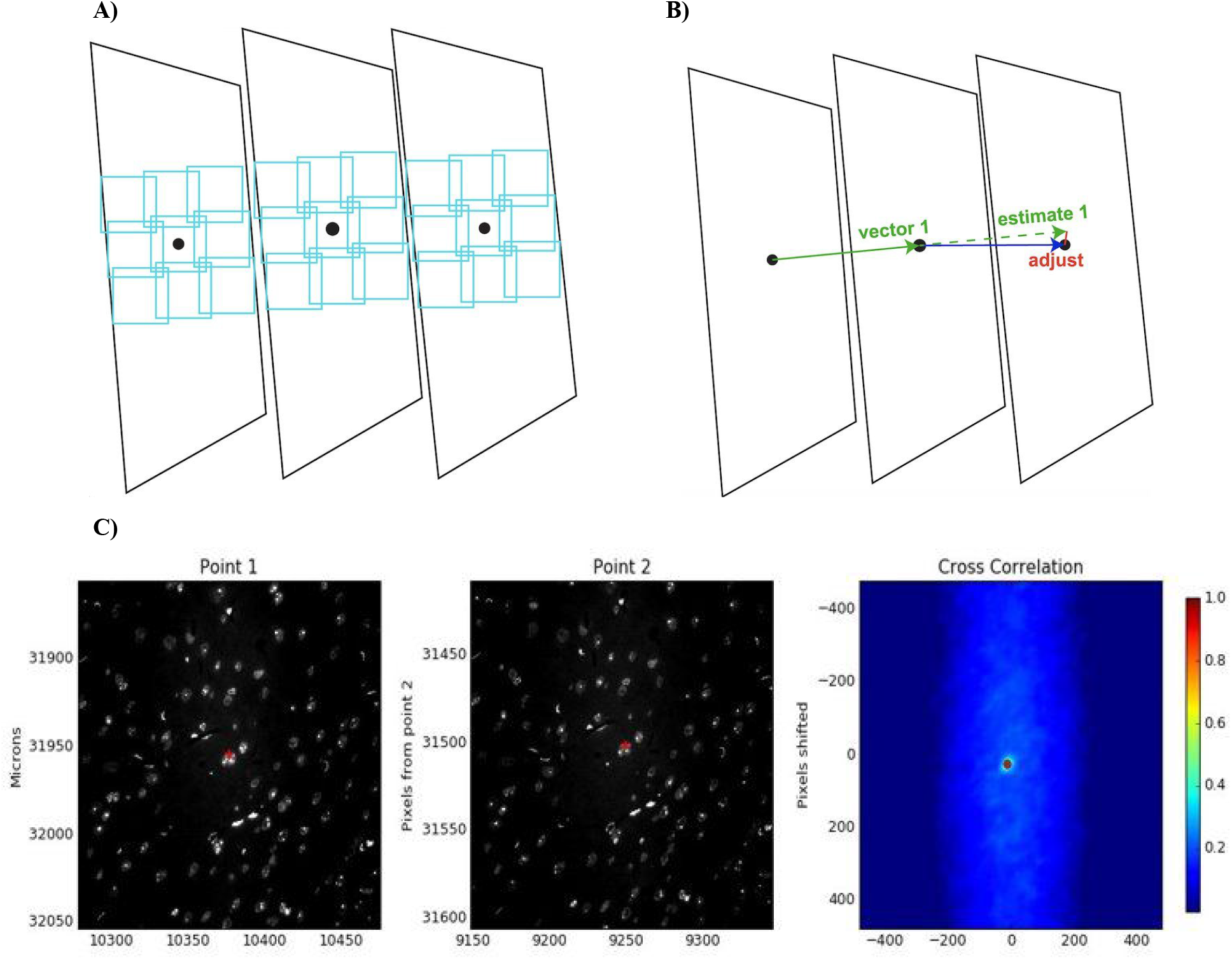
**A) Example layout of consecutive AT sections,** represented as black trapezoids, with a point marking a corresponding feature within a region of interest, individual tiles of which are shown in cyan, on each section. B) The position of a given feature within a sample varies linearly from one section to another, and images of the same feature within adjacent sections are almost identical given that they are separated by ∼ 100 nm axially. The vector from one section to another, starting and ending on the same feature, provides an accurate way to predict where that given feature will be in following section. This prediction can be refined to the pixel level by running a cross correlation of an image captured at the predicted location with an image captured of the same feature in the previous section, and shifting the result to the correlation maximum. C) Images of the same cluster of nuclei, labeled with DAPI, in adjacent AT sections. For the first section, point one, the position of a feature chosen by the user is shown as a red mark, and the axes of this image is the stage position in microns. The second section point 2, is of the same cluster of nuclei in the following section, here the red mark is positioned on the maximum of the cross correlation between the first image and the second. Finally the plot of the cross correlation between the two images is shown on the right.

Fortunately, there are several physical properties of AT samples that can be utilized, given the right software, to address this mapping problem. First, the lateral dimensions of each section are, excluding minor expansion and compression during the sectioning process, the same. Second, because the individual sections within an array are connected together to form a line, the position of any given feature in the tissue can be easily estimated based upon relative positions of that feature in adjacent sections (Figure 1b). Ribbons can curve over many sections, but in general do so gradually, and this linear local estimate remains close across the whole ribbon. Finally, given the section thickness in AT, 50 - 100 nm, images of one feature within the tissue vary little from one section to the next. While the positions of a given feature may vary over the length of the ribbon relative to the substrate the local estimate can be adjusted using simple computer vision (Figure 1c.)

Taking these advantages into account the mapping problem can be broken down into a series of iterative steps which are easily automated. By identifying a vector which points from a given feature within the array section to the exact same feature on an adjacent section, there exists an accurate linear estimate for the location of that exact feature on the section immediately following the latter. The given feature can then be found to within the accuracy of a pixel in this next section by taking an image of the region at the end of the vector and cross correlating it with the image at the estimated location. AT relies on having images of the same regions of tissue from each section present in the dataset. Having a stored set of positions which can be reused within a multi round imaging experiment is necessary for AT data collection, particularly for high throughput experiments. By repeating this process for each pair of sections within an array, this set of points can be collected and used as the basis for creating a tiled acquisition sequence on an AT sample.

MosaicPlanner sets up a user interface to drive this process of linear estimation and refinement in an automated, but also interactive mode that allows the user to intervene and monitor the process. Previous work has utilized this same method, but it was applied to an offline analysis of 10x overview images [*Micheva & Smith 2007, Micheva et al. 2010*] and gave no opportunity for user interaction or correction. MosaicPlanner can both acquire and analyze data, and so it is no longer necessary to acquire 10x overview images, and the mapping process can be driven dynamically. To this end, it should be recognized that there are many procedural differences in conducting AT imaging experiments, especially in regard to sample preparation [*Smith, S. J, 2018*]. MosaicPlanner is most effective when samples are prepared with arrays attached to a coverslip, which in turn is firmly attached to a rigid substrate, reducing the tendency for the coverslip to drift when removed and replaced from a glass slide. MosaicPlanner provides the framework for visualizing individual AT sections, identifying regions of interest, and automating the setup of a reusable imaging plan for an entire array.

## Overview

MosaicPlanner is python based GUI program designed to automate the acquisition of multiplexed light microscopy AT datasets. MosaicPlanner uses the Micro-Manager API to interface with hardware which makes it widely generalizable across many imaging platforms and hardware configurations. Figure 2 depicts a block diagram illustrating both the Micro-Manager hardware groups necessary for MosaicPlanner to interface with the microscope, and examples of hardware within each category which MosaicPlanner has successfully operated.

Data acquisition in MosaicPlanner relies on having a series of points located on precisely the same feature on each section throughout sample. These points, called Slice Positions, provide the framework for capturing tiled images of each section, and ensure that image data acquired covers the same area of tissue throughout the length of the array. The software is designed in such a way that imaging one sample across multiple staining rounds is very simple and robust. The user can set up a series of positions over regions of interest, which are can be saved using stage coordinates, and can then be recalled by the program and reused as needed.

**Figure 2.**
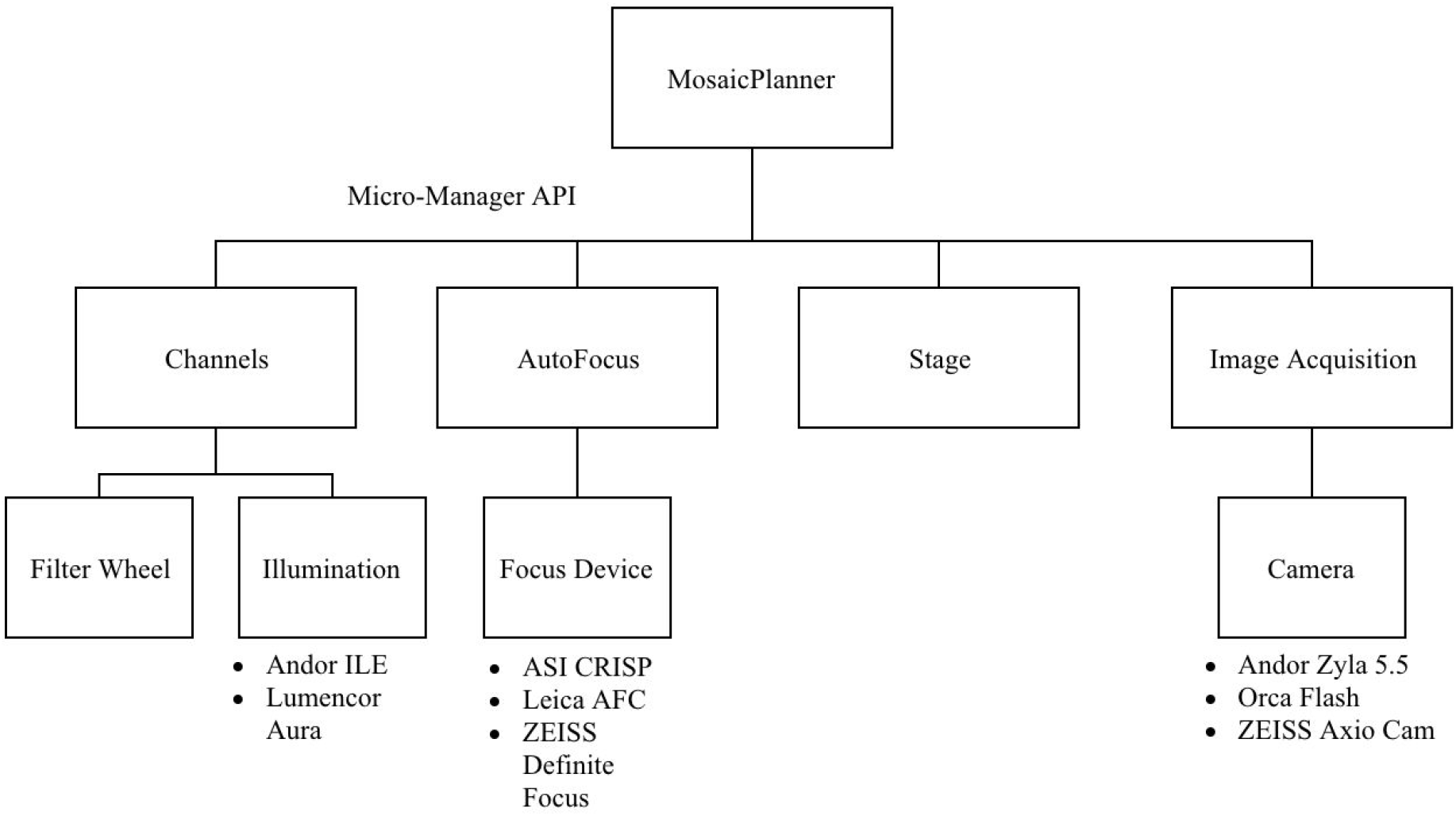
**Software hierarchy within MosaicPlanner.** By using the Micro-Manager API, MosaicPlanner can accommodate many different hardware platforms. The Micro-Manager configuration for the hardware must include a channels group containing an illumination source and filter wheels if present. In addition MosaicPlanner requires a defined autofocus mechanism, stage device, and camera.

## Workflow

During startup the user is prompted to enter in information about the sample they are imaging. MosaicPlanner takes these inputs and uses them to orchestrate a file path system that stores both the data acquired during the mapping stage and the data acquired during the experiment, in a human readable, and machine readable manner. Data storage in MosaicPlanner is discussed later.

## Mapping

Once roughly oriented on the first section, the user can return to the main mapping screen. The user can use the snap button to take a picture at the current location, or utilize tools to snap single images or small mosaic of images at locations the user clicks on the mapping screen. MosaicPlanner will move the stage to that location, perform a focus operation, either software or hardware based depending on capabilities, and acquire an image. That image is then displayed on the mapping screen in its appropriate spatial position, using the stage coordinate space. Using these tools the user can quickly outline the extent of a single section and location a feature of interest within the tissue. Once the user has found a feature of interest they can place a Slice Position on that feature using the add Slice Position tool. Each slice position is used as the center of an M x N matrix of coordinates where each entry represents the center of a frame to be imaged.

Once this process is repeated on the adjacent section, and a slice position is placed on the corresponding feature, the position of that feature within the 3rd section can be identified automatically with the step tool, see supplemental material. MosaicPlanner does so by taking the vector between the first two positions, denoted P1 and P2, moving that distance and direction beyond P2 and snapping an image. MosaicPlanner then takes the central 200×200 pixel cutout of that image, centered on the estimated point, and runs a cross correlation between this new image and the same size cutout centered about P2. MosaicPlanner will add a new slice position on the xy stage coordinates of this maximum. Finally, this new slice position is denoted P2, while the former is changed to P1. This process can be iterated over to automate the setup of a mosaic over a sample, done using the fast forward tool. The fast forward tool will repeat the step process as long as the correlation maximum is above a certain threshold, defined in the software’s configuration. It is helpful if the initial two slice positions are centered on a well defined feature like a nucleus, if using nuclear stains, or a blood vessel, so that refinements of the position list on subsequent rounds of imaging are straight forward. If for whatever reason, the cross correlation falls below the specified threshold, MosaicPlanner will stop this automated process and give control back to the user to adjust the added slice position as needed.

Once the mapping process is completed MosaicPlanner stores each slice position in an object, called a position list, which can be saved and reloaded along with the images taken during mapping. MosaicPlanner position lists also store an orientation, which can be calculated automatically using the rotate tool, which uses the tangent to ribbon’s position list to estimate each section’s orientation. However, MosaicPlanner also allows the user to manually override this orientation for specific sections. The section’s orientation is then used to optimize the layout of the mosaic, given the constraint that individual camera frames can only be taken in a fixed orientation. The current layout is painted for the user so they can evaluate where data will be acquired relative to features of the section. The lateral coordinates of each slice position is individually adjustable as well. Because MosaicPlanner stores the orientation of each section, these lateral adjustments can be made in the relative orientation of each section. On a curved ribbon, moving up the section might be up and to the right in stage coordinates on one section, but up and to the left on another. This feature is invaluable when attempting to optimize the placement of a mosaic to cover a region of interest in all sections. Finally the number of frames can be adjusted by changing the mosaicX and mosaicY parameters, located on the toolbar, as well as the overlap of each frame.

If additional imaging rounds are necessary, the images acquired during the mapping stage can be reloaded into MosaicPlanner. In addition the mosaic settings, called position lists, can be saved as xy-stage coordinates and reloaded. This ensures that each additional dataset is identical, and images precisely the same regions of tissue.

## Acquisition

Once the user has set up their imaging plan, and adjusted the imaging parameters appropriately, data acquisition begins by clicking the start button. Mosaic Planner will move the stage such that the objective is positioned in the top left frame of the first Slice Position. At each frame MosaicPlanner will snap an image in each channel, and place each image in a data queue. The stage will then iterate through each frame in the slice position in a serpentine manner, repeating this process at each slice position. The user can abort acquisition by hitting cancel, and, if using a hardware autofocus mechanism, MosaicPlanner will stop acquisition if focus is lost.
An overview of the entire process is shown in a flow diagram in (Figure 3).

**Figure 3.**
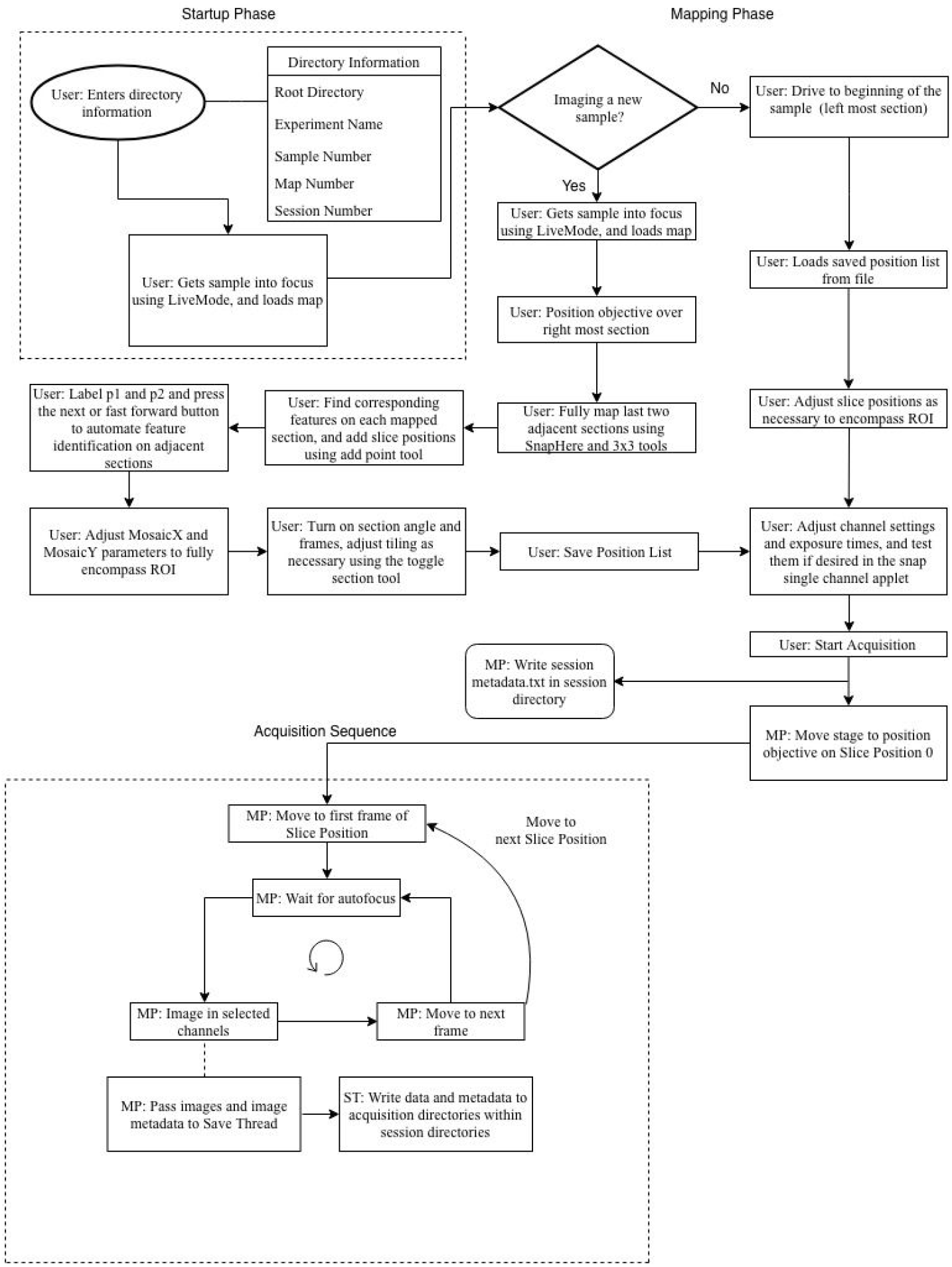
**Flow chart depicting key steps in setup and imaging AT datasets using MosaicPlanner.** The process is broken up into three major phases.1) The startup phase where users enter information about their sample which then allows MosaicPlanner to create data output directories. 2) The mapping phases where users locate their slice positions, set up their tiling sequence, and adjust imaging parameters 3) Acquisition Phase MosaicPlanner automates the acquisition sequence by running through each tile in a repeated pattern

## Retake dialog

While hardware autofocus mechanisms are quite reliable they are not infallible. For a technique like AT, which involves tissue sectioned at ∼100 nm, the stage of the microscope must be precisely coplanar with the focal plane of the objective. Based upon a 1.4 NA 63x objective with an axial PSF of +/− 500 nm the tolerable degree of tilt within a 200×200 um^2 field of view is ∼1 millirad for there to be no noticeable focal gradient in the image. Further, reflection based autofocus can be misdirected if anything abnormal, say a fluorescent piece of dust or bubble in the immersion oil, is present within the field of view. For these reasons MosaicPlanner comes equipped with a mechanism for reviewing and retaking frames which are out of focus.

A focal score is calculated using a Laplacian filter for every frame, and saved in the metadata. In the retake dialog, located under the imaging settings tab on the UI, each frame is plotted by its stage coordinates, and colored based on its focus score. The dialog provides the user with the ability to review their data in a sequential manner, and if necessary refocus and retake data on individual frames.

The retake dialogue offers the user control over the microscope hardware while allowing them to review data from their most recent project. The user can toggle between frames using the w,s,a, and d keys on the keyboard, moving between sections and right and left between individual frames. If the user desires to retake a frame, they can move the objective there, and focus the image using a software autofocus function. This software autofocus function works by using a laplacian filter on a z-stack of images. This is also available to the user to aid them during mapping. The original data from this frame will be moved into a different directory within the project, and replaced in the appropriate channel folders with the newly acquired images.

## Data Storage

The major advantage of AT as a technique is its capability for multiplexing. However this capability is not limitless within any one round of staining. Not all proteins of interest have primary antibodies that are compatible with one another within the same incubation. Further, there is a limit in the number of conjugated secondaries that can be applied due to both excitation and emission spectral overlap. Often this means that for any one AT experiment there needs to be multiple rounds of staining and imaging. In order for data taken from multiple imaging sessions to be useable it not only needs to cover precisely the same areas of tissue, it also needs to be organized in a consistent and machine readable manner, so that downstream image processing steps can interact with the data easily. Unstructured image files can quickly become the source of inconsistencies that make systematic processing of datasets painful. MosaicPlanner saves data organized by an experiment title, sample number, and imaging round; the software saves the last used input, and recalls these settings either during startup or when a new project is started. This process makes it easy for the user to either start a new imaging round of the same sample, or ensure that data from multiple sequential samples are stored in the same place. By saving the map images and position lists, the user only needs to map each sample once, and after reloading the images, adjust each position list slightly to correct for small variations in position when a sample is removed and replaced.

Metadata is saved in an open, transparent machine and human readable format for both the overall experiment, each session directory, and each individual frame. Open human and machine readable metadata reduces barriers for both humans and computer programs to begin interacting with the data. This naming convention makes the data easily navigable and is stereotyped regardless of imaging round. Given that the data is organized in a stereotyped way, image processing scripts based on this organization can be easily implemented. Further, the way in which each individual frame is named running stitching and registration algorithms become much simpler, since the inputs can easily be iterated over.

## Performance Improvements

Having a software platform for AT data allowed us to explore alterations to the acquisition process and optimize the throughput for AT acquisition. General purpose acquisition software must allow the user to configure a virtually unbounded set of possible acquisition routines in order to cover the range of use cases most desirable for a particular application. Too often this results in an interface that is either overly complicated, acquisition routines that are not optimized for the application, or sometimes both.

Overall acquisition times scale linearly and unavoidably with a number of factors including: the surface area of the tissue, number of channels, exposure times, and the length of the sample. However, the acquisition breaks down into the repetition of a single sequence that can be optimized (Figure 3). How much time each of those steps takes depends on several factors in both hardware and software that allowed for more effective optimization with full control over the data acquisition process.

A reflection based autofocus mechanism can reduce the length of an experiment drastically compared to an imaging based focus method (Figure 4), which requires real time data analysis, and additional images to be acquired. However, these devices can be configured and used in a variety of contexts. For example, one could turn the autofocus on for only a brief period of time to recenter the Z position of the microscope at the sample, but then turn it off again. This is in fact usually necessary to take a set of serial images in Z, or to account for any chromatic aberrations. However, for AT applications where only a single plane is necessary, leaving the autofocus engaged in a continuous feedback loop, and simply monitoring when the feedback loop had resolved was the most efficient way to use of autofocus.

**Figure 4.**
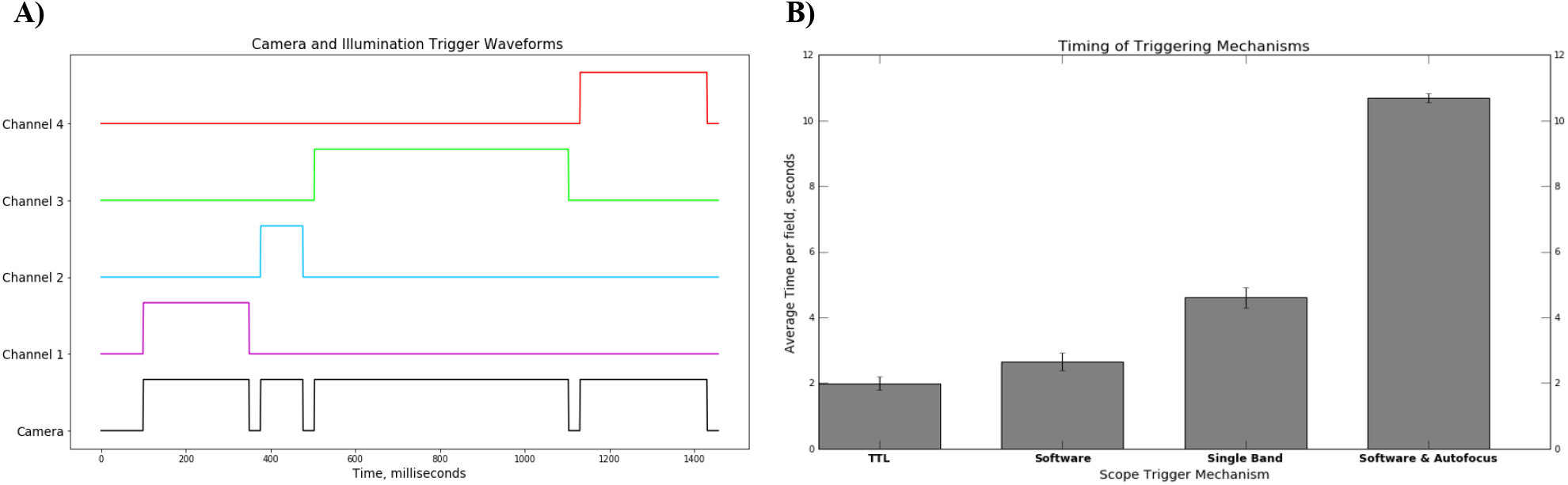
**A)Example waveforms relayed from the Arduino controllers to the camera and laser engine.** The camera shutter is triggered on each individual pulse by one Arduino, while individual illumination shutters are turned on and off by a second Arduino, whose digital state directly controls individual pins for each channel on the laser engine. Most modern scientific cameras have a pixel readout time of ∼25 ms, the ‘down time’ between each pulse must be longer than that. Above is an example using 27 ms between pulses and typical exposure times for each illumination. B) Average times of different triggering mechanisms in MosaicPlanner. Averages are calculated using the difference in total acquisition times over 100 fields of view. All four conditions were taken using the same sample with the same regions of interest and the same exposure times for four channels. A) TTL triggering of the lasers and camera on each frame, while using a quad band dichroic/emission cube, gives us an average time of 1.99 seconds per frame, which can be further improved upon with shorter exposure times and a lower lock threshold for the hardware autofocus. Compared with the average time of software triggering using the same reflection based autofocus and filters, but without hardware triggering over the same fields the result was 2.65 seconds per frame. Average time of acquisition per frame while using a mechanized filter turret which switches positions for each individual exposure yielded 4.61 seconds per frame. The last condition was using an image based autofocus routine on each frame before acquisition, giving 10.7 seconds per frame, increasing the time per frame to over five times that of TTL triggered acquisition.

After stage movement and autofocus, the time to acquire multiple spectral channels is a function of the time it takes to setup the microscope for each channel, by changing illumination, filter wheels and exposure times on the camera, plus the exposure time itself. In practice, the amount of setup overhead, including the amount of time move a conventional filter turret (250-350 ms), the time it takes to configure a camera with a new exposure time (75 ms), and the time to turn on and off a light source or shutter (50-100 ms) can be equal to or longer than the exposure time of a single channel.

There are now many commercial solid state LED and laser illumination devices suitable for widefield fluorescence microscopy. Using narrow band excitation with the appropriate wavelengths can enable the user to forgo using any excitation filter in the light path. Further, by choosing the right fluorophores a multi-band dichroic and emission filter can enable imaging with minimal crosstalk between channels and nullifying the extra time for switching filters entirely. In addition, switching physical filters inevitably introduces small shifts between different spectral images which must be corrected offline.

Importantly, many of these solid state devices are also now switchable via TTL triggering, as are many scientific cameras. Therefore the sequence of different excitation channels and different exposure times can be pre-programmed into a set of TTL pulses delivered to the camera and illumination source in a coordinated fashion, removing most of the overhead involved (Figure 4a). To examine the effect of hardware triggering on acquisition times, two Arduino microcontrollers were used to trigger the camera shutter and laser. The first Arduino is programmed to initiate the sequence of TTL pulses, and relay that sequence to the camera shutter and the second Arduino. This second Arduino is connected to the control pins of each individual laser shutter within the engine. Changing the digital state of this Arduino can turn each shutter on and off in the correct sequence. The cameras that have been used to this end have a pixel readout time of ∼ 25 ms, which we accounted for by using a 27 ms delay between each TTL pulse (Figure 4a).

To demonstrate the practical time savings associated with these improvements, we measured the average acquisition time per field, a single x,y location with four individual channels, across 100 fields. In one configuration, we utilized a quad band filter set and hardware triggering. In a second configuration we used hardware autofocus, and a quad band dichroic, but used software triggering of the camera and illumination. Third we kept the reflection based autofocus but used a single band filter set for each channel, which required a mechanized filter turret and software triggering of the illumination and camera. Finally, we used an image based software autofocus routine, a quad band dichroic, and software triggering of all components. Exposure times, light source and the sample were kept fixed throughout, (Figure 4b).

Hardware triggering the illumination and camera in conjunction with a reflection based autofocus allowed us to image at a rate of just under 2 seconds per xy location, while for the same conditions but software triggering the illumination and camera yielded ∼ 2.65 seconds per field making hardware triggering roughly twenty five percent faster than conventional triggering. Further, while imaging with a mechanised single band filter turret acquisition speeds were slowed to over 4.5 seconds per xy location. Finally, an image based autofocus routine and software triggering of illumination and camera yielded over 10 seconds per xy location.

## Discussion

The first AT datasets were acquired using a fully manual microscope, with the microscopist moving the stage by hand and identifying the corresponding locations in each section by eye, snapping individual images, changing filters by hand, and manually reassembling the resulting image files into coherent volumes. [*Micheva et al. 2010, Micheva & Smith 2007*] Adoption of mechanized microscope components alleviated some of this burden, but setting up position lists remained tedious. Plugins to commercial microscope control software have allowed some automation of position lists, but tend to only work on very well cleaned ribbons with minimal irregularities [*Weiler et al. 2014*]. Other custom software solutions have been described, ZEISS Correlative Array Tomography software for example, but position list creation remains a mostly manual process, that takes up to 20-30 minutes per ribbon. By automating the mapping process we have been able to cut this time in half. Further, MosaicPlanner is set up in such a way that reloading maps of the same sample to set up additional imaging experiments is easy, fast, and organized, which in turn allows for large dataset acquisition.

In the past, AT datasets have covered relatively small surface areas of tissue over short arrays [*Micheva et al. 2010*]. While the technique still provided highly valuable information on morphological structures and the molecular species expressed within them, AT imaging experiments could be time consuming. Many times restrictions were imposed due to practicality, as datasets often took several hours to image, not including time taken for sample preparation or post acquisition image processing. This methodology presents a limitation as biological structures often travel throughout large volumes or cover wide swaths of lateral surface area and thus require a large number of images to examine. This problem is additionally compounded if multiple rounds of staining are required. These limitations can lead to data which only gives a partial view of a potential morphology or region of interest. With the capability of high speed imaging over large surface areas within AT sections, MosaicPlanner offers experimentalists the ability to image the entire tissue volume. After processing and reconstruction they can pick and choose what subregions of their data to examine, and are confronted with the physical limitations of the tissue volume itself, as opposed to those imposed by the imaging.

Through the use of the Micro-Manager API, we have also made this custom software solution available to a wide variety of microscope hardware, which should help lower the barrier of entry for AT. Using a standardized tool for AT acquisition allows image processing pipelines to focus on fewer potential data formats and metadata storage mechanisms. This helps separate the concerns of data acquisition and image processing, allowing innovation in both areas to proceed more independently. Being open source, it is relatively easy for programming savvy experimentalists to integrate novel software or hardware components into the acquisition sequence. The hardware triggering mechanisms described in this paper are one example of a customized hardware system we successfully integrated into this framework.

MosaicPlanner is one example of how a specialized software acquisition system can simplify and facilitate large scale data acquisition. WaferMapper [*Hayworth et al 2014*] is a conceptually similar tool applied to AT acquisition of serial section SEM images that has facilitated high throughput acquisition, though the lack of a standardized SEM microscope API presently limits its broad applicability. SPIM-fluid is an example of an architecturally similar approach applied to light sheet microscopy, where custom software is written on top of the Micro-Manager API. SPIM-fluid carefully coordinates rotational stages, light source and cameras to ease the acquisition of light sheet datasets [*Gualda et al. 2015*].

Given that the structures for data storage are created in a stereotyped and automated manner with some input from the user, MosaicPlanner makes organizing large multi-round AT experiments relatively painless. Data organization can often be an antagonizing process to individuals or teams, and the ability to construct data pathways which are both human and machine readable removes much of the potential for datasets to become disorganized. Thus the software makes it easy for the user to add in an additional staining round or sample into an experiment as these new pathway parameters can just be added into previously existing structures.

AT is a sensitive technique both in sample preparation and image processing. Given the two dimensional nature of the samples, and the infallibility of autofocus mechanisms and stages, we became highly motivated to create a mechanism for reviewing and quality controlling data as it came off the microscope, but before it was processed in any way. This became more important as we began to more frequently work with datasets which required multiple staining rounds or covered a large volume of tissue. The creation of the retake dialog allowed for users to review datasets immediately after acquisition was completed, and touch up any problematic images. This innovation made it that much easier to have datasets which could be easily aligned and registered downstream of the imaging process.

Finally, AT requires reproducibility of imaging conditions, which in turn requires precision and sample visualization. The process of acquiring AT data has been painful in the past, in regards to both the time it takes to set up and acquire the data and the necessary post acquisition image processing. In addition AT datasets can rapidly become large, and thus difficult to organize, which makes processing and interpreting the data that much more diffìcult for the practitioner. The motivation for MosaicPlanner came from these practical factors, and provides a highly flexible platform which lowers the threshold of entry for would be AT experimenters. The software provides a high degree of adaptability for almost any type of imaging condition or issue, and by using the Micro-Manager API, is modular in regards to hardware. MosaicPlanner provides a robust system of both data organization and enables the user to acquire large, multi-round, highly multiplexed AT datasets with relative ease.

## Acknowledgements

We wish to thank the Allen Institute for Brain Science founder, Paul G. Allen, for his vision, encouragement and support.

## Supplemental Material

Icons on the MosaicPlanner toolbar used for mapping and acquisition.

| Icon |  | Icon |  | Icon |  |
| --- | --- | --- | --- | --- | --- |
|  | Adjust map display, select icon then click and drag within display |  | Add Slice Position, select icon then click in display |  | Toggle Position, select icon then click in display, will turn off closest slice position or frame, MP will skip imaging |
|  | Snap image at current stage position, click icon |  | Choose Point 1, select icon then click in display, will set point 1 to be closest slice position |  | Toggle Slice Position Rotation |
|  | Move stage to position, select icon then click in display |  | Choose Point 2, select icon then click in display, will set point 2 to be closest slice position |  | Delete selected Slice Positions |
|  | Move stage and snap at location, select icon then click in display |  | Select Slice Position, select icon then click in display, shift + click can select multiple points, click + 'a' selects all. |  | Display frames on slice positions |
|  | Take 3x3 mosaic on click, select icon then click in display |  | Run Correlation between points 1 & 2, click icon |  | Start Acquisition |
|  | Zoom, select icon, then click and drag within display, will zoom to box |  | Execute step function, estimates where feature will be on next section using vector between points 1 & 2 |  | Execute software autofocus function, click icon |
|  | Crop map display to view all map images, click icon |  | Fast Forward, automates step process, will continue to find feature location until canceled or correlation value is below threshold |  | Open live mode display, click icon |

